# Where are the beachmasters? Unexpectedly weak polygyny among southern elephant seals on a South Shetland Island

**DOI:** 10.1101/2021.05.20.444924

**Authors:** H.J. Nichols, B. Fuchs, A.J Paijmans, G. Lewis, C.A. Bonin, M.E. Goebel, J.I. Hoffman

## Abstract

Intraspecific variation in animal mating systems can have important implications for ecological, evolutionary and demographic processes in wild populations. For example, patterns of mating can impact social structure, dispersal, effective population size and inbreeding. However, few species have been studied in sufficient detail to elucidate mating system plasticity and its dependence on ecological and demographic factors. Southern elephant seals (*Mirounga leonina*) have long been regarded as a textbook example of a polygynous mating system, with dominant ‘beachmaster’ males controlling harems of up to several hundred females. However, behavioural and genetic studies have uncovered appreciable geographic variation in the strength of polygyny among elephant seal populations. We therefore used molecular parentage analysis to investigate patterns of parentage in a small satellite colony of elephant seals at the South Shetland Islands. We hypothesised that dominant males would be able to successfully monopolise the relatively small numbers of females present in the colony, leading to relatively high levels of polygyny. A total of 424 individuals (comprising 33 adult males, 101 adult females and 290 pups) sampled over eight years were genotyped at 20 microsatellites and reproductive success was analysed by genetically assigning parents. Paternity could only be assigned to 31 out of 290 pups (10.7%), despite our panel of genetic markers being highly informative and the genotyping error rate being very low. The strength of inferred polygyny was weak in comparison to previous genetic studies of the same species, with the most successful male fathering only seven pups over the entire course of the study. Our results show that, even in a species long regarded as a model for extreme polygyny, male reproductive skew can vary substantially among populations.

## INTRODUCTION

Understanding mating systems and their evolution is a central goal of behavioural and molecular ecology, animal behaviour and evolutionary biology (Clutton-Brock 1989, Reynolds 1996, Neff & Pitcher 2005, Kempenaers 2008). Mating systems influence a multitude of ecological and evolutionary processes, ranging from social structure, dispersal and gene flow through to the evolution of life-history and sexually selected traits, local adaptation and ultimately speciation (Dieckmann, O’Hara & Weisser 1999, Ross 2001). Understanding mating systems can also have practical implications because of their downstream impacts on inbreeding, effective population size variation and population dynamics, which can influence the ability of populations to respond to challenges such as environmental change (Waser, Austad & Keane 1986, Nunney 1993, Plesnar-Bielak *et al.* 2012).

Most mammalian mating systems are characterised by unequal investment between the sexes in reproduction (Trivers 1972). As a result, males usually compete with one another to mate with as many females as possible, whereas females are often choosy, selecting males who can provide them with either direct or indirect fitness benefits (Fisher 1958, Zahavi 1975, Prokop *et al.* 2012). Sexual selection for indicators of male fitness such as large body size, behavioural dominance and the control of resources results in polygynous mating systems where the variance in reproductive success among males of the same population can be considerable (Clutton-Brock 1988). The extent to which male reproductive success varies among species depends on ecological and phylogenetic factors that determine the distribution of oestrus females over space and time (Emlen & Oring 1977), including the numbers of adult males and females present in breeding groups (Kutsukake & Nunn 2006).

Further complexity arises from the observation that mating systems not only differ among species, but can also vary within species. This is in accordance with theoretical discussions emphasising the importance of ecological and demographic factors shaping the ability of males to gain access to females or the resources required to attract them (Emlen & Oring 1977). Reproductive skew can also be influenced by the costs and benefits that individuals experience of breeding together and the tactics that competitors use to maximise their own reproductive success (Hodge 2009, Clutton-Brock 2016). However, studies of intraspecific variation in mating systems are relatively uncommon and mating systems are often considered as more-or-less fixed attributes of a given species (Gursky-Doyen 2010). Consequently, more studies of intraspecific variation in mating systems are needed both to gain a broader understanding of the magnitude of variation within versus among species and to understand the specific factors responsible for that variation.

The southern elephant seal (*Mirounga leonina*) is a textbook example of a strongly polygynous mammal (Clutton-Brock 2016) that provides an excellent opportunity to investigate variation in animal mating systems. Southern elephant seals spend most of their lives foraging at sea but return to terrestrial haulout sites to breed during the austral summer months. Although foraging often takes place far away from these sites, maternally directed natal philopatry tends to limit the exchange of individuals among breeding colonies (Nichols 2009). Consequently, four main, genetically distinct populations have been recognised: South Georgia in the South Atlantic Ocean, Heard and Kerguelen Islands in the South Indian Ocean, Macquarie Island in the South Pacific Ocean, and the Valdés Peninsula population on the coast of mainland Argentina (Slade *et al.* 1998, Hoelzel, Campagna & Arnbom 2001, McMahon *et al.* 2005). However, less intensively studied breeding populations can also be found on other sub-Antarctic islands, including the South Shetlands (Hunt 1973).

Southern elephant seals exhibit extreme sexual size dimorphism, with males being approximately six times larger than females (González‐Suárez & Cassini 2014). During the breeding season, females congregate on beaches to pup and re-mate, and dominant males (known as a ‘beachmasters’) fight to control harems of females (defined as a group of females with a dominant male in attendance). Behavioural studies at South Georgia (Laws 1956, McCann 1980, Modig 1996), the Falkland Islands (Galimberti, Fabiani & Sanvito 2002), Marion Island (Wilkinson & Van Aarde 1999) and Macquarie Island (Carrick & Ingham 1962) have shown that a handful of the highest ranking males can monopolise harems of many tens to over a thousand breeding females on densely packed beaches. These dominant males tend to be older and larger than subordinate males, who attempt to gain mating opportunities by entering harems to secure ‘sneaky’ matings, intercepting females as they leave the harems to forage at sea (McCann 1981), and potentially by mating with females at sea (De Bruyn *et al.* 2011). Although these alternative mating strategies do appear to have a limited payoff for subordinate males, genetic studies of this species have shown that behavioural observations generally provide a reasonable proxy of male reproductive success, with harem holders typically accounting for a large proportion (up to 90%) of all paternities (Wainstein *et al.* 1997, Hoelzel *et al.* 1999, Fabiani *et al.* 2004). Furthermore, beachmasters have been known to hold harems over multiple consecutive years (Fabiani *et al.* 2004), which could result in the most successful males fathering hundreds of pups over their lifespans.

However, polygyny need not always be this extreme in elephant seals. For example, southern elephant seals on the Valdés Peninsula breed at relatively low density due to the availability of hundreds of kilometres of uninterrupted, open beaches. This results in harems that are on average smaller than those observed at other localities (median = 11 females per harem, range 2–122) within which females are spaced out (e.g. single harems of over 100 females take up at least 12,000 m^2^ of beach), making it difficult for harem holding males to monopolize matings (Baldi *et al.* 1996). For example, dominant males with large harems (over 50 females) have a high number of copulations (37 per 100 hrs), but these harems are difficult to control by a single male and females frequently mate with subordinate males (9 copulations per 100hrs) (Baldi *et al.* 1996). While the degree of polygyny on the Valdés Peninsula is lower than at high density breeding colonies, harem holders still have high reproductive success, achieving 55% of observed copulations and 58% of paternities (Hoelzel *et al*. 1999), while 75% of males achieve no paternities (Wainstein 2000).

The most southerly breeding sites for southern elephant seals are located in the South Orkney Islands and the South Shetland Islands (Laws 1956). These populations could potentially have different patterns of male reproductive skew than previously studied populations for two main reasons. First, at Signy Island in the South Orkneys, around 70 pups per year were recorded as having been born in 4–6 harems (Laws 1956). These small harem sizes will limit the maximal reproductive success of harem holders simply because these males will have access to fewer breeding females. Second, in particularly cold years, breeding takes place on fast ice where space is unrestricted, leading to even greater female dispersion and potentially lower male reproductive skew (Laws 1956). However, genetic studies have not been conducted in these localities, so the realised degree of polygyny is unknown. Conversely, smaller harems might in fact be easier for dominant males to monopolise, allowing beachmasters to attain relatively high reproductive success at low population densities. This was previously shown for a low density colony of Antarctic fur seals in the South Shetlands, where the most successful males surpassed the reproductive success of dominant males at a high density colony in South Georgia (Bonin *et al.* 2014). Genetic studies are therefore required to assess the degree of reproductive skew in lower density colonies of southern elephant seals and to further understand the drivers of variation in polygyny.

Here, we use genetic parentage analysis to investigate the degree of polygyny of southern elephant seals at Half Moon Beach, Cape Shirreff, in the South Shetland Islands (Figure 1). The relatively small Cape Shirreff breeding colony is considered a ‘satellite’ to South Georgia because there is some exchange of individuals between these locations (Boyd, Walker & Poncet 1996). Together with other islands in the archipelago, such as King George Island and Elephant Island, Cape Sherriff may represent a stop-off point for breeding females migrating northwards to larger colonies from southerly pelagic foraging grounds closer to the Antarctic Front, with some females remaining there to breed (Krzemiński 1981, Laws 1994). In warmer, more favourable years, many females breed at Cape Sherriff, whereas in colder, less favourable years, the accumulation of sea ice and snow causes fewer animals to come ashore. Consequently, the size of the breeding population varies appreciably from year to year. Similar to the Valdés Peninsula, the beach at Half Moon Bay represents a large expanse of uninterrupted breeding habitat that is sparsely occupied by southern elephant seals in the breeding season. In most years, there is only one harem, but in years of high pup production there can be as many as five harems.

**Figure 1.**
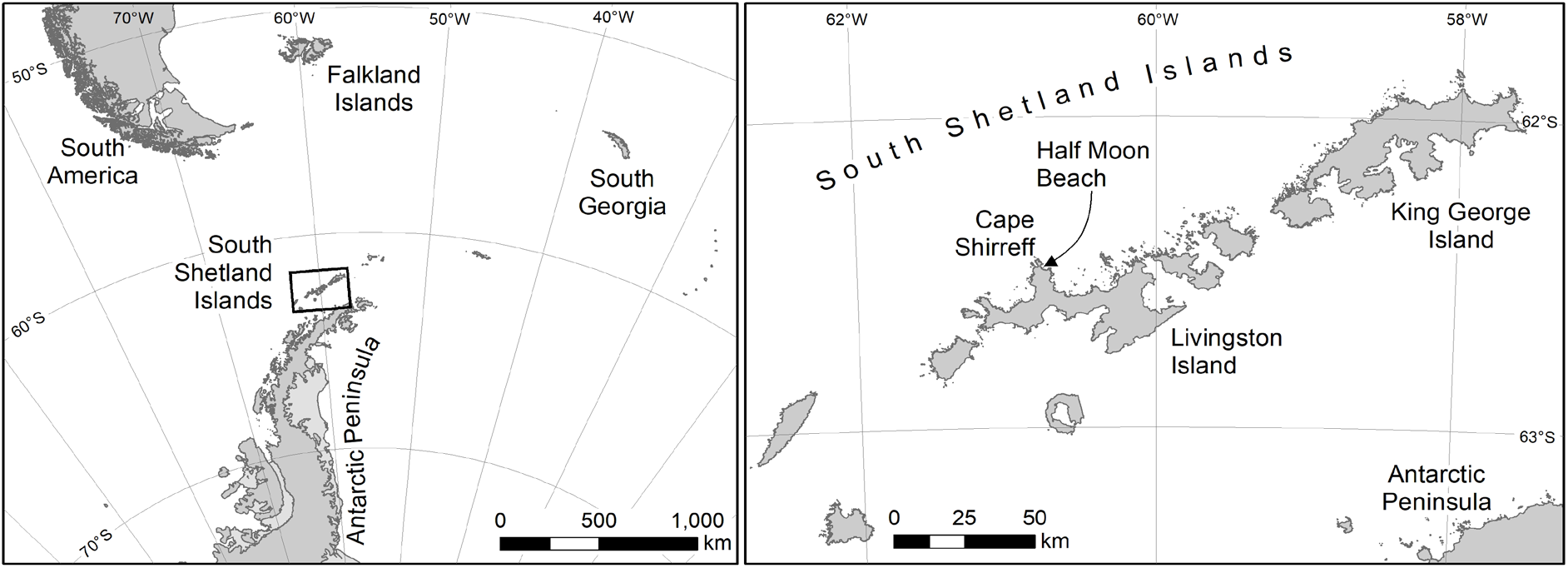
The study location and terrain at Half Moon beach, Cape Shirreff, in the South Shetland Islands.

In order to investigate the strength of polygyny in this breeding colony, we sampled all of the pups born across eight breeding seasons, together with the majority of adult males and a large number of breeding females. To maximise the power to resolve parental relationships, we genotyped all of the individuals at 20 highly variable microsatellites. Wherever possible, maternities were assigned in order to maximise the power of the paternity analysis and to provide a benchmark against which patterns of male reproductive success can be compared. As previous studies of Antarctic fur seals have shown that male reproductive skew is higher at low density (Hoffman, Boyd & Amos 2003, Bonin *et al.* 2014), we hypothesised that the mating system of southern elephant seals at Cape Shirreff would be highly polygynous, with a small proportion of males monopolising reproductive success. Both the degree of reproductive skew between competing males and the size of the harem they compete over are likely to contribute to dominant male reproductive success. We therefore also predicted that the maximum annual reproductive success of harem holders may be lower at Cape Shirreff than at high density breeding sites, due to the smaller sizes of the harems.

## MATERIALS AND METHODS

### Study site and sample collection

This study was conducted at a small breeding colony at Half Moon Beach, Cape Shirreff, South Shetland Islands (62°28’37.4”, 60°46’45.0”, Figure 1). Cape Shirreff is an ice-free peninsula, covering approximately 3.1 km^2^. It is the most northerly point on the north coast of Livingston Island. Half Moon Beach is a horseshoe shaped beach stretching over 1.65 km. It is a wide-open sandy / cobble beach with areas of higher elevation that contain a few large boulders and one long rocky outcrop extending perpendicularly relative to the waterline and located approximately 70 m from the low tide line. Elephant seals tend to concentrate around these areas of the beach, typically in a single harem but sometimes in as many as five, which are separated by as much as 400 m (Figure 2). During the breeding season (October–November) pupping areas are covered with between a few centimetres and two meters of snow. Access to harems often requires navigating a substantial berm of ice and snow at the high tide mark. Due to the remote setting and challenging conditions, researchers typically arrive at Cape Shirreff midway through pupping (mean arrival date 29 October, s.d. = 7.3 days), leading to the inference that the southern elephant seals start hauling out at the breeding site in early October. The mean number of animals present during the breeding season in our study was 106 (s.d. = 37.7; range = 56–158). As in other populations, harems were dominated by a single male, with subordinate males present on the periphery or elsewhere on the beach. Approximately 95% of all adult males, including all harem holders and the majority of peripheral males were sampled together with all of the pups and females that were present on the beach. Individuals were carefully approached at the harems in order to minimise disturbance.

**Figure 2.**
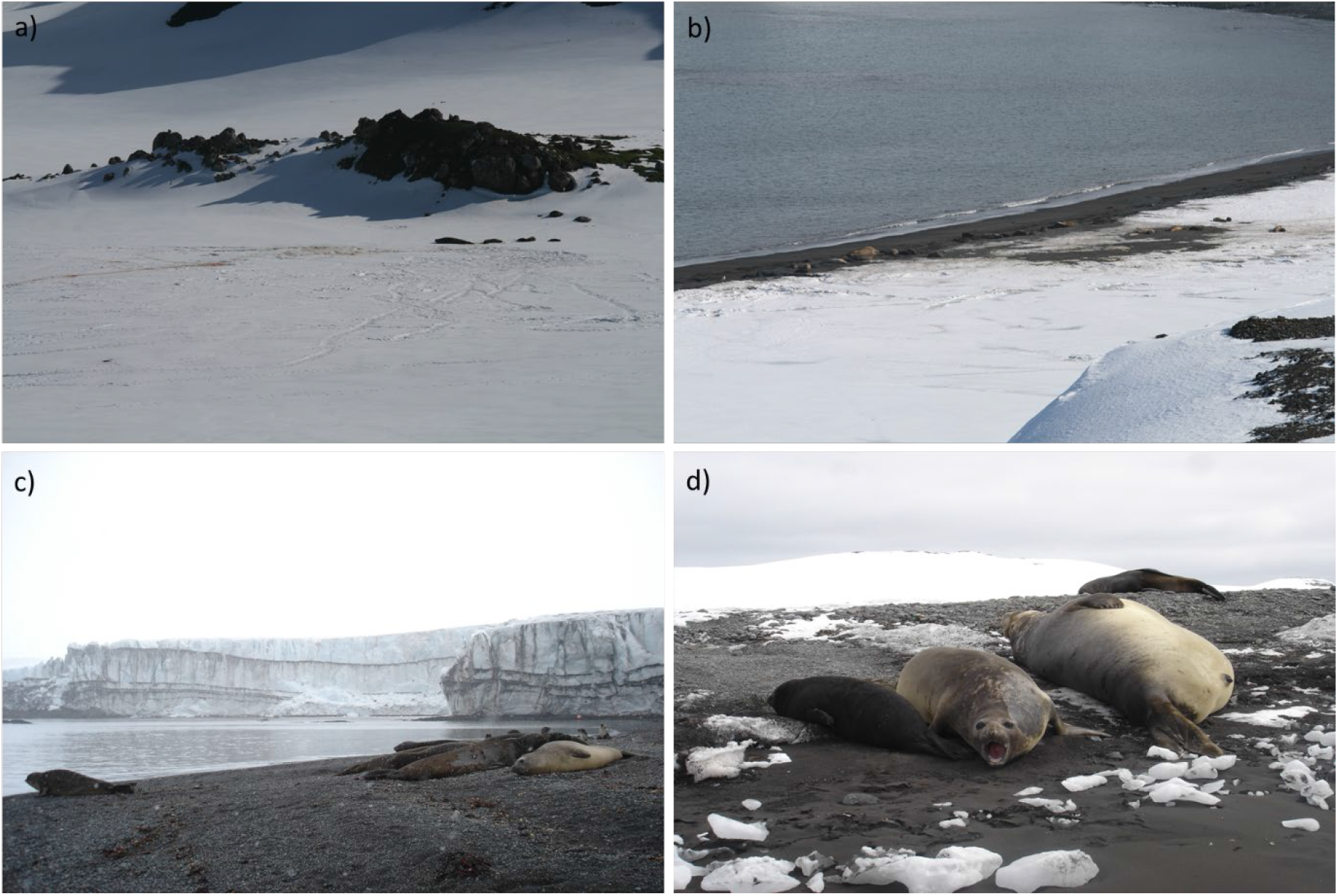
Southern elephant seal harems at Half Moon beach, showing a) a small harem located close to an elevated rocky area that could potentially provide cover for males pursuing alternative mating tactics, b) and c) harems close to the shoreline at high tide, and d) a female (centre) with her pup and a harem holding male.

Fieldwork for this project was conducted during the Austral spring (October to December inclusive) of 2008 to 2016, except for 2010 (Table S1). Samples were collected during 2010 but unfortunately these were lost in transit from Cape Shirreff back to the USA. In most seasons, adults were sampled on the same day, but when the numbers of individuals were large, they were sampled by harem over consecutive days. Adults were sampled from the flanks using a 2 mm sterile, disposable Miltex™ biopsy punch (Fisher Scientific). Pup skin samples were collected from a rear flipper while they were captured for tagging using a tag hole punch or a 2 mm, sterile, disposable Miltex biopsy punch. Skin samples were immediately placed in 95% ETOH and kept frozen (−20°C) until analysis. Sampled adults were marked using hair dye whenever the bulk of sampling efforts spanned more than a day (sampled individuals were otherwise identified by their biopsy punch mark). Marks persisted throughout the season but were lost at the moult so were not informative from one season to the next. Most individuals were sampled within a few days of the first visit to the colony but the site was checked for unsampled individuals thereafter during broad pinniped censuses at Cape Shirreff and / or to retrieve instrumentation.

### Microsatellite genotyping

DNA was extracted using a standard chloroform/isoamylalcohol protocol (Sambrook, Fritsch & Maniatis 1989) and genotyped at 20 polymorphic microsatellites (see Table S1 for details). PCR amplifications were performed in five separate multiplexed reactions using a Qiagen^®^ Multiplex PCR Kit following the manufacturer’s recommendations, except that we used 12 μl reaction volumes to keep the use of reagents to a minimum. The following PCR profile was used: one cycle of 5 min at 94 °C; 24 cycles of 30 s at 94 °C, 90 s at 60 °C, 30 s at 72 °C; and one final cycle of 15 min at 72 °C. Between 12 and 20 positive controls were included on each 96-well PCR plate to facilitate the standardisation of microsatellite scoring across plates. Fluorescently labelled PCR products were resolved by electrophoresis on an ABI 3730xl capillary sequencer (Applied Biosystems, Waltham, MA, USA) and allele sizes were scored using GeneMarker v. 2.6.2 (SoftGenetics, LLC., State College, PA, USA). To maximise genotype quality, we manually inspected all of the traces and corrected any genotype calls where necessary.

### Quantification of the genotyping error rate

Genotyping errors can strongly influence the outcome of genetic parentage analyses (Marshall *et al.* 1998, Hoffman & Amos 2005). We therefore took the precaution of independently re-extracting and re-genotyping a total of 96 randomly selected samples. The resulting duplicate genotypes were then used to calculate the error rate per genotype and per allele for each locus separately and combined over all loci.

### Genetic data analysis

We removed 33 samples that were genotyped at <15 loci and then checked the dataset for duplicate genotypes (representing resampling events) using the R package “poppr” (Kamvar, Tabima & Grünwald 2014). This identified 19 individuals that were sampled two or more times, usually in different years. Excluding these samples resulted in a final dataset of 424 unique individuals comprising 33 adult males, 101 adult females and 290 pups. We calculated the observed and expected heterozygosity of each locus using R package “adegenet” (Jombart 2008). We then tested for deviations from Hardy-Weinberg equilibrium (HWE) based on 10.000 Monte Carlo permutations using the R package “pegas” (Paradis 2010). The resulting *p*-values were corrected using the Bonferroni correction in the p.adjust function of the “stats” R-package. Finally, we tested for population substructure by implementing a principal component analysis (PCA) of the dataset using “adegenet”. Because PCA can be sensitive to missing data, any missing genotypes were imputed and the allele frequencies were transformed by centring and scaling the data. The probability of identity and the exclusion probability were calculated using “GenAIEx” version 6.5 (Peakall & Smouse 2005).

### Parentage analysis

Parentage analysis was conducted for the 290 pups within “COLONY” 2.0.6.6 (Jones & Wang 2010). All sampled adult males (33 individuals) and females (101 individuals) were included as potential parents and we did not specify any known parents or sibships. We set weak priors that the true mother and father were in the candidate lists as 0.5 and 0.2 respectively and specified a polygynous mating system in a diploid species. We used a medium run size, which provides high confidence parentage assignments without exceeding practicable run-times. Parentage assignments were accepted with ≥ 0.95 probability. An advantage of COLONY is that, instead of using pairwise comparisons to assign parentage, it uses a full-pedigree maximum likelihood approach, which considers the likelihood of the entire pedigree structure and allows the simultaneous inference of both parentage and sibship. This allows the program to assign genetically unsampled individuals as parents, which provides additional insights into mating patterns despite the incomplete sampling of parents. Colony has been shown to be highly accurate and is the only program available that can use genetic information to assign offspring to unsampled parents (Harrison *et al*. 2013, Walling *et al*. 2010). Wilcoxon tests were conducted in R base package to investigate differences in reproductive success between different categories of individual.

## RESULTS

Our final dataset for parentage analysis comprised 424 southern elephant seal individuals genotyped at 15–20 microsatellites (for details, see Table 1 and Table S1). The loci carried on average 9.3 alleles and none deviated significantly from Hardy-Weinberg equilibrium after table-wide Bonferroni correction (Table S1). The genotyping error rate, calculated by independently repeat-genotyping 96 samples, was low at 0.003 (0.3%) per allele or 0.006 (0.6%) per genotype. The probability of identity was 1.13 × 10^−21^ and the exclusion probability was 1.

**Table 1.** Details of the numbers of unique southern elephant seal individuals (on the basis of multilocus microsatellite genotypes) including a summary of the parentage assignments, best configuration fathers and breeding sex ratios (BSRs) calculated based on the number of males and females sampled on the beach and using parentage data from pups born at the study site.

| Year | Number of<br>adult males | Number of adult<br>females | Number of pups | Number of<br>mothers assigned | Number of fathers<br>assigned | Number of best<br>configuration<br>fathers | BSR for<br>sampled<br>individuals | BSR for best<br>configuration<br>parents |
| --- | --- | --- | --- | --- | --- | --- | --- | --- |
| 2008 | 8 | 17 | 36 | 19 | 1 | 18 | 2.13 | 1.06 |
| 2009 | 6 | 15 | 19 | 14 | 1 | 12 | 2.5 | 1.17 |
| 2011 | 6 | 9 | 57 | 10 | 1 | 8 | 1.5 | 1.25 |
| 2012 | 3 | 6 | 26 | 10 | 3 | 7 | 2 | 1.43 |
| 2013 | 4 | 17 | 33 | 18 | 3 | 16 | 4.25 | 1.13 |
| 2014 | 2 | 2 | 14 | 4 | 3 | 3 | 1 | 1.33 |
| 2015 | 1 | 33 | 54 | 23 | 2 | 23 | 33 | 1 |
| 2016 | 3 | 2 | 51 | 8 | 3 | 7 | 0.67 | 1.14 |
| Mean | 4.13 | 12.63 | 36.25 | 13.25 | 2.13 | 11.75 | 5.88 | 1.19 |

### Parentage analysis

COLONY assigned mothers to 108 pups and fathers to 31 pups, of which 18 had both parents assigned. Because so few pups were assigned fathers, we used two complementary approaches to analyse patterns of paternity. First, we focused on the small number of genotyped fathers assigned to 31 pups, and second, we analysed the *best configuration fathers* inferred by COLONY. Best configuration fathers include genotyped fathers together with “hypothetical genetic fathers” inferred using maximum likelihood to explain the offspring genotypes. Using the offspring genotype and the genotype of the assigned mother together with data on allele frequencies, COLONY is able to assign unsampled fathers to pups, including assigning the same unsampled father to full or paternal half siblings. Best configuration parentage assignments therefore allow us to investigate likely patterns of paternity, even when many fathers remain unsampled. When analysing best configuration fathers, we conservatively restricted our analysis to the 108 pups for which maternity could be assigned, thereby excluding pups with neither parent sampled.

At least one pup was assigned to 75% of genotyped females (*n* = 76 mothers) and 36% of genotyped males (*n* = 12 fathers) over the course of the study (individuals with at least one pup assigned are subsequently referred to as mothers and fathers). Mothers almost exclusively had a single pup in any given year, with the exception of three pairs of twins (Figure 3a). Annual reproductive success was significantly higher in fathers than mothers, whether limiting the analysis to genotyped fathers (Wilcoxon-test: W = 543, *p* < 0.0001) or best configuration fathers (Wilcoxon-test: W = 548, *p* = 0.002). Nevertheless, male reproductive skew was modest within years, with the majority of fathers only being assigned one pup (56% of genotyped fathers, Figure 3b; and 87% of best configuration fathers, Figure 3c).

**Figure 3.**
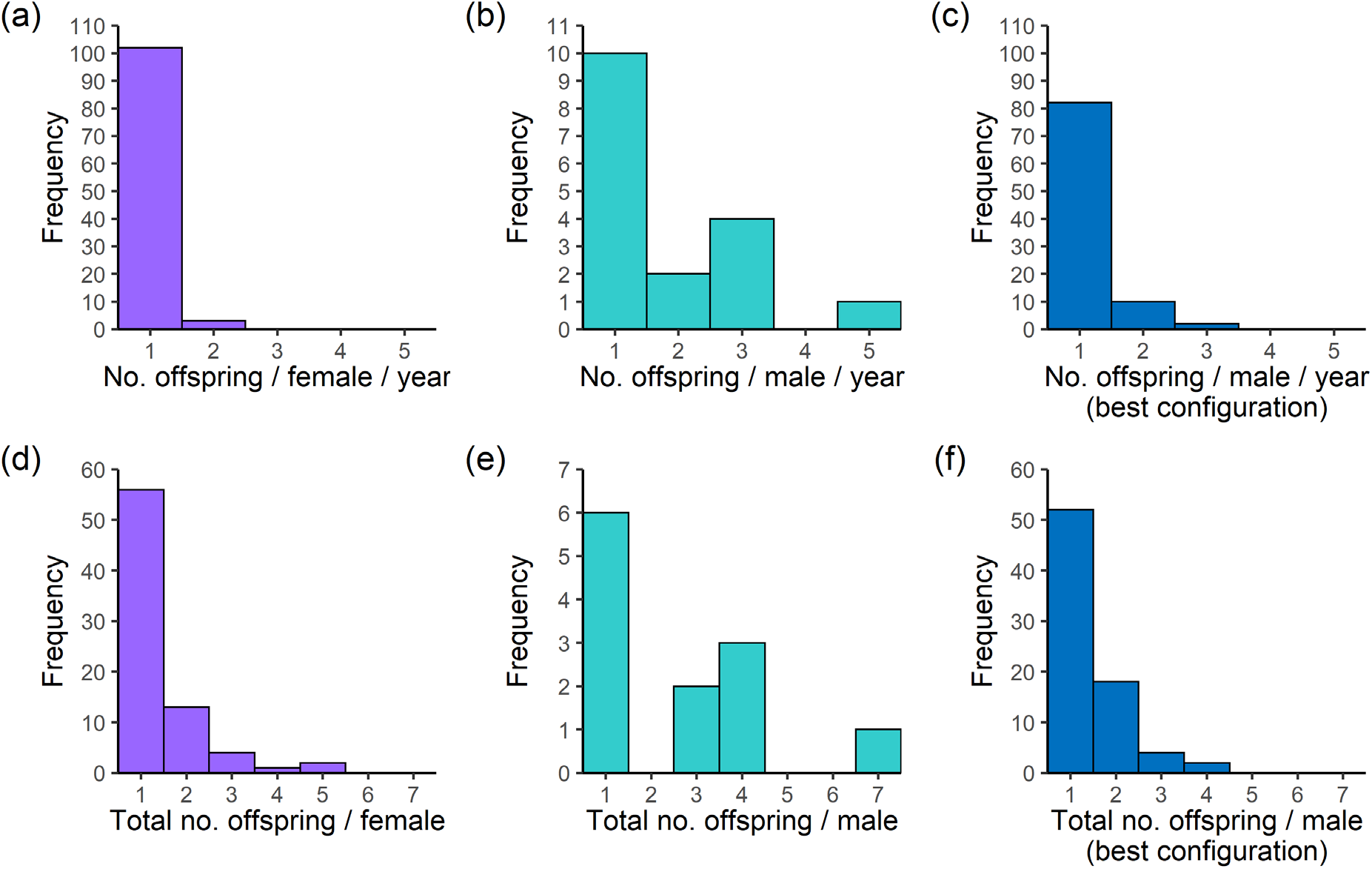
Reproductive skew in southern elephant seals inferred from parentage assignments (panels a,b,d,e) as well as from the best configuration fathers inferred by COLONY (panels c,f). Note that the latter was conservatively inferred from pups that were assigned maternity to a genotyped female, thereby excluding pups with neither parent sampled. The number of offspring assigned to genotyped males was, in some cases, larger than the number of offspring assigned to best configuration fathers because ~ 40% of paternities were assigned to pups that were not assigned to a genotyped mother. Panels (a-c): number of offspring assigned per year of study (where the individual was assigned at least one offspring in the given year); panels (d-f): number of offspring assigned combining all years.

Total reproductive success over the eight years of the study showed a qualitatively similar pattern, with genotyped fathers again producing more offspring on average than genotyped mothers (Wilcoxon-test: W = 680, *p* = 0.008) although the difference was small, with females producing a maximum of five offspring (Figure 3d) and males a maximum of seven (Figure 3e). There was no difference in the total number of pups assigned to genotyped mothers and best configuration fathers (Wilcoxon-test: W = 3012, *p* = 0.57, Figure 3f).

The majority of individuals (50% of genotyped fathers, 77% of best configuration fathers and 75% of genotyped mothers) only produced offspring in a single breeding season. Low reproductive skew could therefore result from high turnover of breeders, with many females only visiting the beach once, and hence mating with unsampled males in other locations the previous year. To investigate patterns of reproductive success in individuals that consistently bred at the study site, we identified a subset of ‘core’ genotyped individuals that were associated with the beach across multiple years based on genetic recaptures and parentage assignments (Figure 4). These comprised 11 males (33% of sampled males), 24 females (24% of sampled females) and the 67 pups that were genetically assigned offspring of these core adults (at least one parent was a core adult). In no year did a single male monopolise all of the offspring produced by core individuals (Table S2); the most successful male in each year fathered between 17% and 60% (mean = 32%) of these pups, equivalent to a mean of 2.6 pups per year (range = 1–5). Furthermore, PCA did not identify any obvious genetic differences between core and transient individuals (Figure 5), or between pups that were and were not assigned a father (Figure 6), implying that they most likely originate from a single panmictic population.

**Figure 4.**
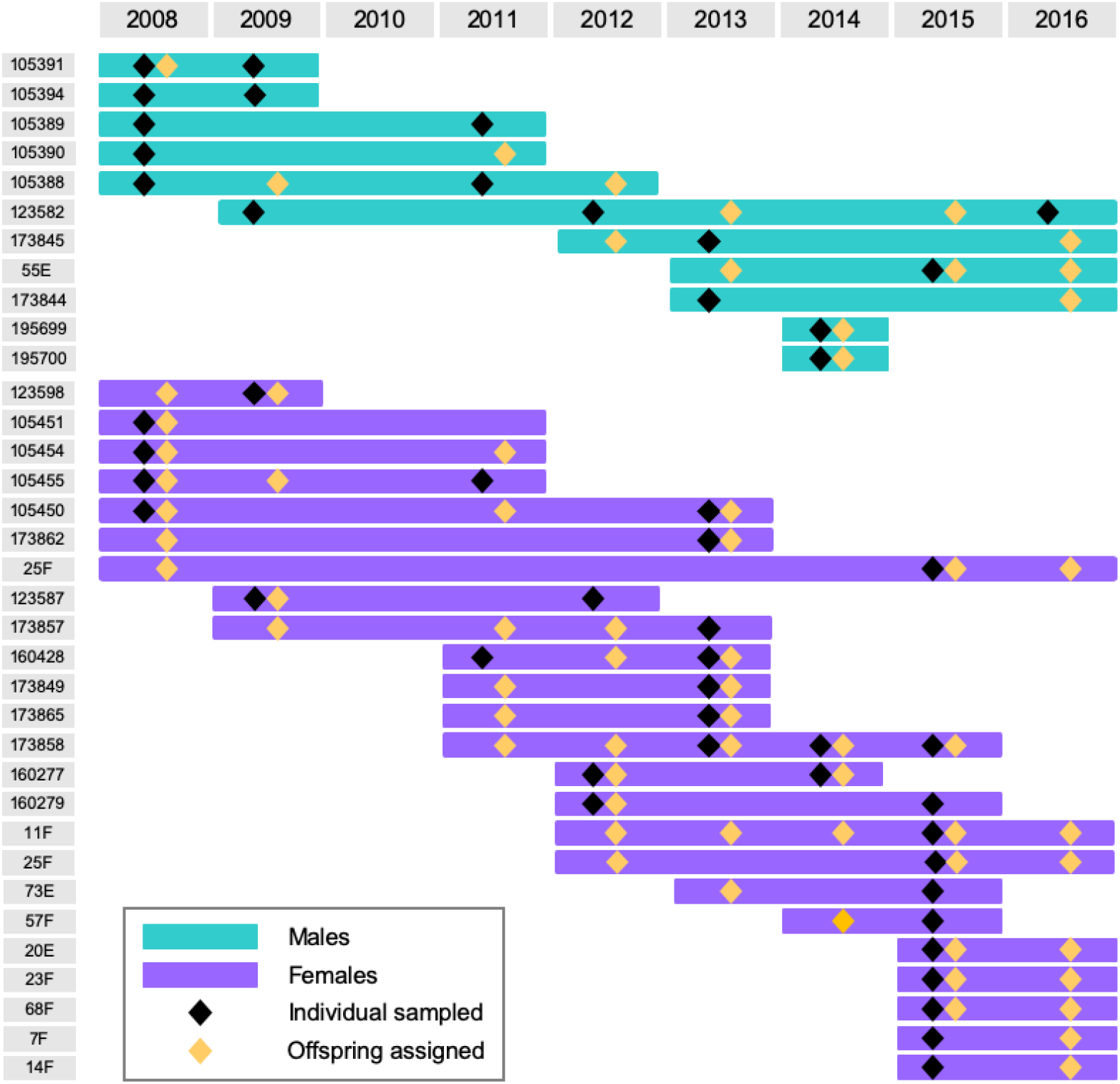
Breeding histories of ‘core’ elephant seal individuals inferred from genetic recaptures (black diamonds) and parentage assignments (yellow diamonds). Males are shown above in green and females below in purple.

**Figure 5.**
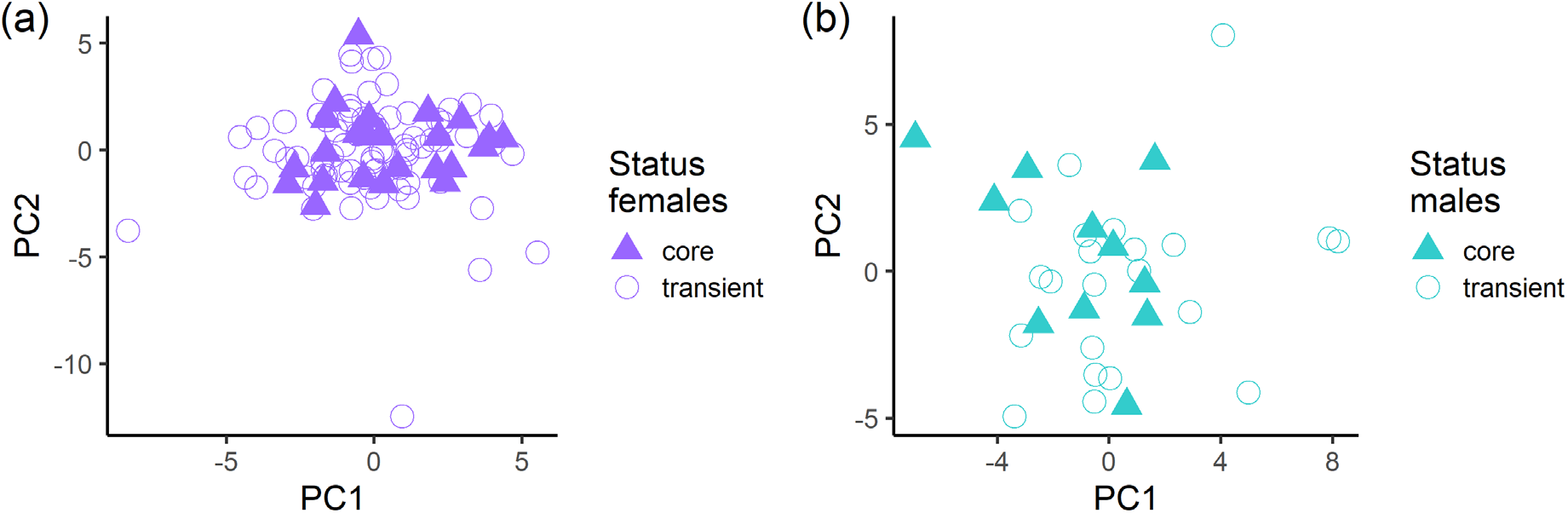
Results of PCAs of (a) breeding females; and (b) breeding males. The points represent individual variation in principal components one and two. Symbol-colour combinations distinguish between ‘core’ and ‘transient’ breeders as defined in the main text.

**Figure 6.**
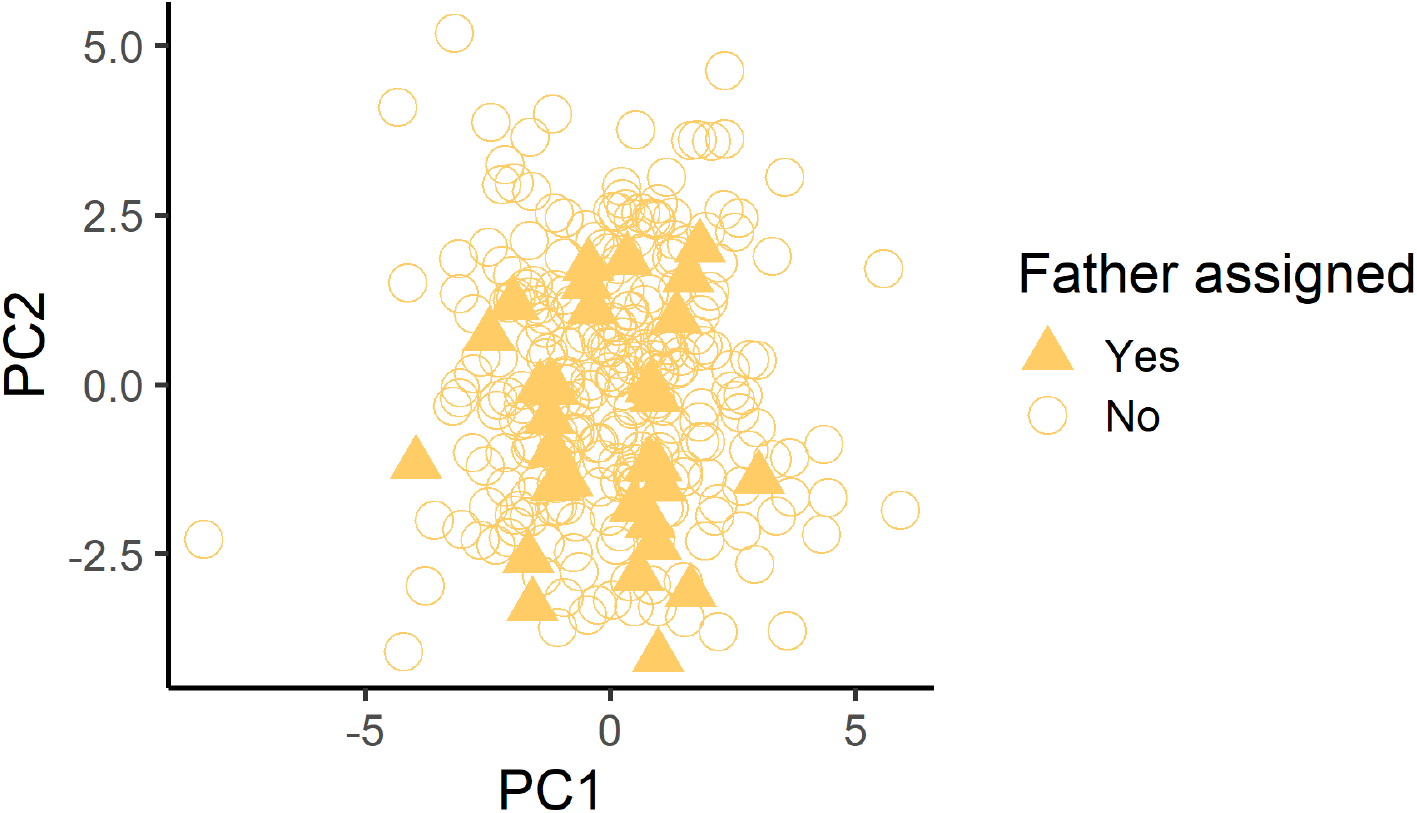
Results of PCA of 290 southern elephant seal pups. The points represent individual variation in principal components one and two. Symbol-fill combinations distinguish between pups assigned paternity and pups not assigned paternity.

Finally, although core breeders returned to the colony across multiple years, they were not always present every year. Consequently, many comparisons between candidate males and offspring will involve males who were not present in the colony during the year a given pup was conceived, while some pups may also have been conceived elsewhere if their mother was absent from the colony in the conception year. We therefore restricted our analysis to include only comparisons involving mothers who were known to be present in the colony during the year of conception. Although our sample size was reduced to only 20 pups, we did not find evidence for strong polygyny (Table S3). Specifically, no single male fathered more than two offspring per year, reproduction was monopolised by a single individual in only one year, and many of the pups (35%) were fathered by unsampled males.

## DISCUSSION

Southern elephant seals have long been regarded as a classic example of extreme polygyny (Clutton-Brock 2016), with ‘beachmaster’ males monopolising large harems of breeding females (Carrick, Csordas & Ingham 1962) and fathering up to 90% of all offspring (Wainstein et al. 1997, Hoelzel *et al*. 1999, Fabiani *et al*. 2004). However, previous studies of this species have uncovered appreciable variation in the environmental potential for polygyny. We therefore investigated patterns of parentage at a southern elephant seal breeding colony in the South Shetlands, where low densities and high rates of turnover among animals may lead to different outcomes in terms of male reproductive success. We found that polygyny was relatively weak at Cape Shirreff, where paternity could only be assigned to around ten percent of pups and reproductive skew was only slightly stronger in males than in females. We discuss these findings in the context of reproductive skew in harem-holding species and mating system flexibility, as well as in light of specific features of the focal population.

### Paternity assignment rate and strength of inferred polygyny

We were only able to assign paternity to 31 out of 290 pups (10.7%). This low rate of paternity assignment contrasts with previous molecular genetic studies of southern elephant seals at the Valdés Peninsula, South Georgia and the Falkland Islands, where 58%, 74% and 90% of paternities respectively were attributed to harem holders (Wainstein *et al*. 1997, Hoelzel *et al*. 1999, Fabiani *et al*. 2004). We also found that the strength of polygyny at the South Shetlands was much weaker than observed in similar studies of elephant seals at other localities. Specifically, the most successful males in our study were assigned up to five offspring in a single season and up to seven pups across all seasons combined. By comparison, harem holders at the Falkland Islands were assigned between 25 and 32 paternities in a single year (Fabiani et al. 2004).

These findings are surprising because in theory small harems should be easier for behaviourally dominant males to monopolise. Indeed, across species, larger harems have been shown to have greater rates of extra pair paternity, with the dominant males of larger harems losing out on a larger proportion of paternities (Clutton-Brock & Isvaran 2006, Isvaran & Clutton-Brock 2007), and the same may also be true within species. In the Antarctic fur seal, another pinniped that breeds in both low and high density colonies (Meise *et al.* 2016), dominant males appear to achieve greater reproductive success at low density (Hoffman, Boyd & Amos 2003, Bonin *et al.* 2014) and pairs of individuals repeatedly re-mate across years, implying a relatively static mating system in which territorial males are more successful at monopolising access to breeding females. So why do southern elephant seals at Cape Shirreff deviate from the expected pattern? We can think of a number of possible explanations, broadly classified into (i) methodological aspects such as the quality of the genetic data and the completeness of sampling; (ii) demographic factors such as small population size and high breeding female turnover; and (iii) alternative reproductive strategies.

### Methodological aspects

The importance of methodology cannot be understated in molecular genetic parentage studies. For example, even a relatively modest genotyping error rate of 1% per allele can result in over 20% of true fathers being excluded from paternity when around ten microsatellites are used (Hoffman & Amos 2005). We guarded against this possibility by implementing strict quality control measures including carefully standardising allele scoring across plates through the inclusion of multiple positive controls, manually checking all traces and independently re-extracting and blind genotyping almost a quarter of the animals selected at random. As the resulting error rate estimate was only a fraction of a percent, we conclude that genotyping errors are very unlikely to explain the paternity shortfall.

Previous studies of pinnipeds have also emphasised the importance of the completeness of male sampling. For example, the rate of paternity assignment in Antarctic fur seals varies from year to year in relation to the percentage of sampled males (Hoffman, Boyd & Amos 2003), while disparities between the outcomes of independent parentage studies of grey seals (Worthington Wilmer *et al.* 1999, Twiss *et al.* 2006) have been attributed to mismatches in the spatial and temporal coverage of male sampling. Male sampling biases could potentially have a bearing on our results given that logistical constraints prevented us from sampling throughout the entire duration of the breeding season at Cape Shirreff. Specifically, the annual sampling and marking of adults could not usually be initiated before the last week of October. We know from studies of other elephant seal colonies that female haulout activity usually peaks sometime between the 2^nd^ and the 25^th^ of October, and that the males usually arrive around a week earlier (Galimberti and Boitani 1999). Any males that were exclusively present in the earlier part of the season will therefore not have been sampled. However, our analysis of the best configuration fathers revealed a similar overall pattern to the parentage assignments and did not provide any indications of the presence of a small number of disproportionately successful males. It is therefore unlikely that our results can be explained by a failure to sample a small number of highly successful males. If anything, the magnitude of polygyny inferred from the best configuration fathers appeared to be slightly lower than the magnitude of polygyny inferred from paternity assignments. However, parentage analysis was performed on the full dataset while the best configuration fathers were only inferred for a subset of pups with known maternity.

### Demographic factors

Our study colony of southern elephant seals at Cape Shirreff colony differs markedly from other sites where elephant seals have previously been studied. First, it is relatively small, with a mean of 36 females pupping each year in one to five harems. By contrast, population sizes and harem sizes are much larger at Sea Lion Island (mean females per harem = 47.7, range = 18–91; Fabiani *et al*. 2004), Peninsula Valdes (mean females per harem = 65.3, range = 30–119; Campagna, Lewis & Baldi 1993), South Georgia (mean females per harem = 74.2, range = 6–232; McCann 1980), Isles Kerguelen (mean females per harem = 102, range = 5–1350; Van Aarde 1980) and Macquarie Island (mean females per harem = 277, up to 1000; Carrick, Csordas & Ingham 1962). The smaller harems present at Cape Shirreff may therefore limit the number of females that harem-holding males can monopolise, leading to reduced reproductive skew among males. The Cape Shirreff colony also fluctuates in size across years, with pup production varying from 14 in 2014 to 57 in 2011, whereas northerly breeding populations appear to be more stable. These fluctuations appear to be related to environmental conditions, whereby larger numbers of females breed at Cape Sherriff in warmer years, but in colder years the accumulation of sea ice and snow prevents many animals from coming ashore. As a result, around three quarters of our study females only pupped at Cape Shirreff in a single year, whereas 40– 60% of females are known to return to breeding colonies at the Peninsula Valdes over consecutive years (Hoelzel *et al.* 1999).

This high rate of turnover of breeding females may impact our results because many pups are likely to have been conceived at other breeding colonies, thus diluting the perceived reproductive success of the harem holders at Half Moon beach. To investigate how this might impact our results, we restricted our analysis to pups born to females that were known to be present in the colony during the year of conception. Although our sample size was substantially reduced, the overall pattern of paternity assignment was qualitatively similar and again we did not find any evidence of high male reproductive skew. Additionally, pups fathered at different colonies might be expected to carry different genetic signatures given that the four main global populations of southern elephant seals show pronounced genetic differentiation (Slade *et al.* 1998, Hoelzel, Campagna & Arnbom 2001). However, we could not find any obvious genetic differences between pups with known fathers and pups that were not assigned paternity. This implies that the majority of pups were probably not conceived at distant localities, although we cannot discount the possibility that females may mate at closer, less genetically differentiated sites such as elsewhere in the South Shetland Islands or at South Georgia.

### Alternative mating strategies

Alternatively, mating might take place in or around the study colony but involve alternative reproductive strategies. For example, the unique topography of Half Moon Beach may facilitate alternative male mating strategies on land. The beach covers large areas of sandy substrate at low tide, but the animals mostly congregate around an elevated rocky area located approximately 70m from the low tide line. Consequently, peripheral males only have to cross a short section of empty beach in order to access an area of high topographic complexity (rocky outcrops) that may provide sufficient cover for sneaky copulations. High complexity breeding sites with gullies and dunes appear to diminish the reproductive success of northern elephant seal (*M. angustirostris*) harem holding males because, even though these features concentrate females who might otherwise sparsely occupy a beach, they also provide cover for more agile peripheral males attempting to infiltrate harems to copulate (Hoelzel *et al.* 1999).

Alternatively, the large expanse of unoccupied breeding habitat at Half Moon Beach might facilitate alternative female mating strategies by making it challenging for harem holding males to patrol against other males that would have ample opportunities for aquatic mating. Aquatic mating is relatively common in true seals and has previously been advocated as a possible explanation for the inability of molecular studies to assign paternities in grey seals (Worthington Wilmer *et al.* 1999) and California sea lions (Flatz *et al.* 2012). In addition, De Bruyn *et al.* (2011) argued that aquatic mating may be an important alternative female mating strategy in southern elephant seals given that breeding females often skip coming ashore yet still conceive pups. Given the lack of behavioural data in our study and our inability to sample throughout the entire breeding season (researchers arrive at Cape Shirreff midway the pupping season and miss early arriving animals), we would caution against drawing premature conclusions. Nevertheless, our findings clearly point towards much lower levels of polygyny than observed at other localities (Wainstein *et al.* 1997, Hoelzel *et al.* 1999, Fabiani *et al.* 2004) and highlight the need for future studies focusing on both male and female reproductive strategies (De Bruyn *et al.* 2011).

### Broader perspectives

Harem-based mating systems such as those of pinnipeds exemplify the extreme variation in reproductive success that can occur when a subset of the most behaviourally dominant males monopolise access to breeding females or the resources on which they depend (Clutton-Brock 2016). However, a growing number of genetic studies have brought into focus the importance of alternative mating tactics such as aquatic mating, which can result in lower than expected levels of polygyny. Among polygynous pinnipeds, for example, female grey seals appear to exhibit a combination of partner fidelity (Amos *et al.* 1995) and mate choice directed towards unrelated partners (Amos, Worthington Wilmer & Kokko 2001), while overall, the proportion of offspring that can be assigned paternity is much lower than expected from observations of animals on land, implying that many females may mate at sea (Worthington Wilmer *et al.* 1999). Similarly, in Antarctic fur seals, California sea lions and New Zealand fur seals, female mobility (Hoffman *et al.* 2007, Flatz *et al.* 2012) and alternative male mating strategies (Caudron *et al.* 2010) appear to undermine polygyny. Our study contributes towards this growing body of research by showing that the variation in polygyny within a pinniped species can be greater than previously envisioned. Furthermore, our observation of weak polygyny in a low density southern elephant seal breeding colony is at odds with our original expectations based on Antarctic fur seals, where reproductive skew appears to be higher at low density (Hoffman, Boyd & Amos 2003, Bonin *et al*. 2014). This suggests that findings from one species cannot be readily extrapolated to another, and that specific features of breeding colonies such as topology, female mobility and the amount of exchange of individuals with other colonies, are likely to be important drivers of intraspecific variation in polygyny.

### Conclusion

Studies of intraspecific variation in mating systems are still relatively uncommon and mating systems are often considered to be more or less fixed attributes of a given species (Gursky-Doyen 2010). However, mating systems are the outcome of the reproductive strategies of members of a species rather than evolved characteristics of a species (Clutton-Brock 1989). Hence, they often vary in accordance with the prevailing social and ecological conditions (Carranza, Hidalgo De Trucios & Ena 1989, Bradley *et al.* 2005, Gursky-Doyen 2010, Maher & Burger 2011, Jin *et al.* 2016). This is exactly what we found in southern elephant seals, where pups born at Half Moon Beach in the South Shetland Islands appear to be fathered by a large number of males that each sire between one and a handful of offspring. Several factors may contribute toward this pattern, including incomplete male sampling and breeding female turnover, but these do not appear sufficient on their own to explain the relatively low reproductive success of harem holders, hinting at the possible involvement of alternative mating strategies.

## ACKNOWLEDGEMENTS

All sampling was conducted in accordance with Marine Mammal Protection Act Permit Nos. 16472-01 and 774-1847-04 granted by the Office of Protected Resources, National Marine Fisheries Service, the Antarctic Conservation Act Permit Nos. 2012-005 and 2008-008. The protocols used in this study were also reviewed and approved by the U.S. National Marine Fisheries Service Southwest and Pacific Islands Region’s Institutional Animal Use and Care Committee (SWR/PIR IACUC approval documents: #SWPI2011-02, #SWPI2014-03). We thank the following people for field assistance: Scott Freeman, Ryan Burner, Ray Buchheit, Kevin Pietrzak, Jay Wright, Melany Zimmerman, McKenzie Mudge, Wiley Archibald, Whitney Taylor, Sam Woodman, Raul Vasquez del Mercado, David Vejar, Jessica Senzer, Michelle Goh, Matt Klosterman, Naira de Gracia, Drs. Nicola Pussini, Whitney Taylor and Douglas Krause. A special thanks to the directors of the US-AMLR Program, Drs. Rennie Holt and George Watters. This work could not have been done without the support of the captain and crew of the Laurence M Gould and the National Science Foundation. We are also grateful to Laura Gerrish from the Mapping and Geographic Information Centre of the British Antarctic Survey for preparing the map.

## AUTHOR’S CONTRIBUTIONS

M.E.G. and J.I.H. designed the study, M.E.G. and C.B. performed the fieldwork, J.I.H and B.F. performed the genotyping, H.J.N., A.J.P., G.L. and J.I.H. analysed the data, and J.I.H., H.J.N. and C.B. wrote and finalised the manuscript with input from all of the authors.

## DATA ARCHIVING

Data will be uploaded to a suitable online archive (such as Dryad) on article acceptance.

