## Supplementary Information for "Where are the beachmasters? Unexpectedly weak polygyny among southern elephant seals on a South Shetland Island"

**Table S1.** Details of the microsatellite loci used in this study, including literature sources, polymorphism characteristics in 424 unique individuals and genotyping error rates. $H_e$, expected heterozygosity; $H_o$, observed heterozygosity, HWE $p$-value, uncorrected exact Hardy-Weinberg equilibrium test results based on 10,000 Monte Carlo permutations.

| Locus | Reference | Multiplex | Number of alleles | $H_e$ | $H_o$ | HWE $p$-value | Bonferroni corrected $p$-value | Probability of identity | Exclusion probability | Error rate per allele | Error rate per genotype |
|---|---|---|---|---|---|---|---|---|---|---|---|
| Hg3.6 | (Allen et al. 1995) | 1 | 9 | 0.80 | 0.80 | 0.13 | 1.00 | 0.064 | 0.917 | 0 | 0 |
| Pv9 | (Allen et al. 1995) | 1 | 5 | 0.54 | 0.56 | 0.37 | 1.00 | 0.261 | 0.772 | 0.0026 | 0.0053 |
| Mang27 | (Sanvito et al. 2013) | 1 | 9 | 0.74 | 0.73 | 0.70 | 1.00 | 0.091 | 0.917 | 0.0108 | 0.0215 |
| Mang01 | (Sanvito et al. 2013) | 1 | 5 | 0.72 | 0.69 | 0.68 | 1.00 | 0.124 | 0.772 | 0 | 0 |
| Mang09 | (Sanvito et al. 2013) | 1 | 21 | 0.89 | 0.88 | 0.15 | 1.00 | 0.020 | 0.983 | 0.0027 | 0.0053 |
| Ag-9 | (Hoffman et al. 2008) | 2 | 7 | 0.58 | 0.58 | 0.53 | 1.00 | 0.261 | 0.871 | 0 | 0 |
| ZcCgDh7tg | (Hernandez-Velazquez et al. 2005) | 2 | 10 | 0.81 | 0.81 | 0.49 | 1.00 | 0.062 | 0.932 | 0 | 0 |

| | | | | | | | | | | | |
|---|---|---|---|---|---|---|---|---|---|---|---|
| ZcCgDh1.8 | (Hernandez-Velazquez et al. 2005) | 2 | 12 | 0.87 | 0.89 | 0.90 | 1.00 | 0.031 | 0.951 | 0 | 0 |
| M11a | (Hoelzel et al. 1999) | 2 | 6 | 0.78 | 0.72 | 0.03 | 0.61 | 0.083 | 0.831 | 0 | 0 |
| Hl-8 | (Davis et al. 2002) | 2 | 10 | 0.74 | 0.74 | 1.00 | 1.00 | 0.109 | 0.932 | 0 | 0 |
| Lw-16 | (Davis et al. 2002) | 3 | 7 | 0.77 | 0.75 | 0.10 | 1.00 | 0.088 | 0.871 | 0.0053 | 0.107 |
| Hg8.10 | (Allen et al. 1995) | 3 | 7 | 0.75 | 0.75 | 0.28 | 1.00 | 0.101 | 0.871 | 0 | 0 |
| Lw-20 | (Davis et al. 2002) | 3 | 12 | 0.83 | 0.77 | 0.20 | 1.00 | 0.046 | 0.951 | 0.0027 | 0.0054 |
| ZcwA12 | (Hoffman et al. 2007) | 4 | 10 | 0.76 | 0.77 | 0.91 | 1.00 | 0.095 | 0.932 | 0 | 0 |
| ZcwE03 | (Wolf et al. 2005) | 4 | 5 | 0.58 | 0.59 | 0.26 | 1.00 | 0.221 | 0.772 | 0 | 0 |
| ZcCgDh4.7 | (Hernandez-Velazquez et al. 2005) | 4 | 13 | 0.85 | 0.86 | 0.68 | 1.00 | 0.042 | 0.958 | 0.0058 | 0.0117 |

| | | | | | | | | | | | |
|---|---|---|---|---|---|---|---|---|---|---|---|
| Pvc1 | (Goodman 1997) | 4 | 10 | 0.63 | 0.62 | 0.64 | 1.00 | 0.161 | 0.932 | 0 | 0 |
| Mang44 | (Sanvito et al. 2013) | 4 | 15 | 0.86 | 0.86 | 0.45 | 1.00 | 0.035 | 0.968 | 0 | 0 |
| Lw-8 | (Davis et al. 2002) | 5 | 10 | 0.39 | 0.37 | 0.01 | 0.19 | 0.390 | 0.932 | 0.0380 | 0.0598 |
| Mang35 | (Sanvito et al. 2013) | 5 | 7 | 0.78 | 0.79 | 0.89 | 1.00 | 0.082 | 0.871 | 0 | 0 |
| Total | – | – | 9.5 | 0.73 | 0.73 | – | – | 1.13 x $10^{-21}$ | 1 | 0.0035 | 0.0061 |

**Table S2**. Details of the parentage of ‘core’ pups. Pups were assigned as core if either of their (genotyped) parents had produced pups on the beach in multiple years, or had been sampled in multiple years but only produced one pup.

| Year | Number of best configuration fathers | Number of pups | Maximum number of pups fathered by a male | Proportion of pups fathered by the most successful male |
|---|---|---|---|---|
| 2008 | 6 | 6 | 1 | 0.17 |
| 2009 | 2 | 5 | 3 | 0.60 |
| 2011 | 6 | 8 | 2 | 0.25 |
| 2012 | 6 | 9 | 3 | 0.33 |
| 2013 | 7 | 11 | 5 | 0.45 |
| 2014 | 5 | 6 | 2 | 0.33 |
| 2015 | 8 | 9 | 2 | 0.22 |
| 2016 | 9 | 13 | 3 | 0.23 |

**Table S3.** Results of the parentage analysis restricted to comparisons involving genotyped males and mothers who were both present in the colony during the year of pup conception. The total sample size of pups available for this analysis was 20. Colony best configuration results showed that the six pups with unsampled fathers in 2015 were fathered by five different males (one male fathered two of the pups).

| Conception year | Number of pups | Number of pups fathered by the most successful male sampled in the year of conception | Number of pups fathered by males sampled in a year other than the year of conception | Number of pups with unsampled fathers | Number of males sampled on the beach in the year of conception that were assigned offspring |
|---|---|---|---|---|---|
| 2008 | 2 | 1 | 0 | 1 | 1 |
| 2011 | 3 | 2 | 1 | 0 | 1 |
| 2012 | 3 | 1 | 2 | 0 | 1 |
| 2013 | 2 | 2 | 0 | 0 | 1 |
| 2014 | 2 | 0 | 2 | 0 | 0 |
| 2015 | 8 | 2 | 0 | 6 | 1 |
